# Quantifying the net effect of biodiversity on stability

**DOI:** 10.1101/2025.04.24.650410

**Authors:** Charlotte Kunze, Dominik Bahlburg, Maren Striebel, Toni Schott, Ian Donohue, Helmut Hillebrand

**Author notes:** **Statement of authorship:** C.K., H.H., I.D., and M.S. designed the research. C.K. performed the experiment and analysed the data with assistance from T.S. and D.B. D.B. implemented the model. C.K. wrote the first draft of the manuscript with substantial input from H.H., and I.D. All authors contributed to the drafts. **Data and code availability statement:** All code used for the analysis of the experimental data is publicly available on GitHub https://github.com/charlyknz/MicrocosmExp22, data of the experiemt are archived on Figshare: https://doi.org/10.6084/m9.figshare.25568490.v3. Code for the simulations is publicly available on Github: https://github.com/charlyknz/NBES the data are archived on Zenodo: https://doi.org/10.5281/zenodo.15274625.

## Abstract

Understanding the relationship between biodiversity and both the functioning and stability of ecosystems has been a central focus of ecologists for decades. A step-change in our understanding of the biodiversity–ecosystem functioning relationship was enabled by explicit measurement of the additional functioning provided by biodiversity through comparing expected and observed yields in multi-species communities. However, we lack an equivalent measure for stability. Here, we quantify the net biodiversity effect on stability using model simulations and a microcosm experiment that exposed different phytoplankton species and their combinations to temperature increases and fluctuations. As an emergent property of communities, stability frequently exceeded the expected stability of the combined component species, leading to a net biodiversity effect on stability analogous to the effect on functioning. In our simulations, these effects depended on the strength of competitive interactions as well as species composition and their thermal niche. Experimentally, the stabilising effect of diversity was, however, non-linear, greatest for two-species combinations, and varied with both community composition and disturbance regime. Quantifying the net biodiversity effect on stability advances our mechanistic understanding of the biodiversity–stability relationship, and provides crucial information to support ecosystem management and conservation.

## Introduction

Exploring the intricate interplay between biodiversity and ecosystem stability has been a focal point of ecological research for decades, with a multitude of studies exploring the factors that underlie the stability of ecosystems^1^. Ecological stability comprises a family of measures that together encapsulate the dynamics of the system and its response to disturbance, and include the ability of a system to withstand a disturbance (resistance), recover from it (recovery), and capture how it varies over time (temporal stability)^2,3^.

Recent advances in modelling and synthesis have enhanced our understanding of how stability might be influenced by the diversity of species in a system^4–9^, demonstrating that there is no universal diversity–stability relationship. Rather, individual dimensions of stability differ in their relationships with biodiversity, depending on the nature of environmental change^8,10,11^. For example, increasing species richness decreased resistance to experimental drought in plant communities^10^. Similarly, increasing species richness in microbial communities has been shown to decrease resistance to experimental warming but increase temporal stability^8^. This richness-dependent effect on resistance indicates that species-rich systems may be more likely to lose growth potential under disturbance if species growth is negatively correlated with disturbance tolerance^10,12^. By contrast, diversity has been found to stabilise community dynamics over time (that is, reduce temporal variability) when ecosystems are exposed to environmental variability^6,13^, even though individual populations within diverse communities might exhibit enhanced fluctuations^14^.

Thus, a diverse community can buffer against environmental fluctuations because different species respond asynchronously to disturbances^15,16^. Understanding to what extent stability emerges from biodiversity requires knowledge about how population stability relates to community stability in the context of disturbances. Distinguishing the role of individual species responses from the aggregated effects of interspecific interactions is therefore crucial to predict biodiversity-stability relationships in the context of environmental change^17^.

The related question of how biodiversity affects ecosystem functioning has benefited immensely from such a decomposition, which has become a fundamental element of understanding net biodiversity effects^18,19^. By quantifying expected functions based on species-specific performance in monoculture and comparing these expectations with observed rates in multi-species communities, Loreau & Hector^19^ highlight the dual importance of both species diversity (via complementarity) and species identity (via selection effects) in ecosystem functioning^19^. What is currently lacking is, however, an equivalent metric to quantify the additional stability provided by biodiversity in the context of environmental change.

Here, we extend Loreau & Hector’s^19^ approach for quantifying the net biodiversity effect on ecosystem functioning to assess the added stability provided by biodiversity and, therefore, quantify the net biodiversity effect on functional stability (NBES) — that is, the stability of an aggregate community function such as biomass^3^. The NBES is determined from the difference between observed and expected stability and can be quantified for any metric of stability that is measured by comparing a treatment response to a control (Box 1).

Here, we combine model simulations and a microcosm experiment to determine (i) whether the NBES is consistently positive and (ii) increases with species richness, leading to greater stability in more diverse communities, (iii) how species thermal optima influence the NBES under different temperature regimes, and (iv) how the NBES is influenced by competition strength. We address these questions using the Overall Ecological Vulnerability (OEV)^20^ index as our focal measure of stability, as it combines multiple dimensions of post-disturbance stability into a single measure of instability and therefore allows a direct comparison across different disturbance regimes^20^. We quantify the NBES in model simulations using a temperature-dependent version of the Lotka-Volterra competition model, followed by a microcosm experiment, where we manipulated phytoplankton species richness and temperature disturbance regimes (that is, diurnal temperature fluctuations, a gradual temperature increase, and their combination; see *Methods*). We measured community- and species-specific responses to disturbance and determined the NBES for each species combination in both simulations and the experiment, to test ***H1,*** that the net effect of biodiversity on stability is positive (that is, NBES > 0), as the observed stability in species combinations is higher than expected from species in isolation; ***H2*** that NBES increases with increasing species richness, ***H3*** that NBES varies among species combinations and temperature treatments due to differences in species thermal optima ^21–23^, and ***H4,*** that the magnitude of the NBES depends on the strength of competitive interactions, as interactions directly shape species responses to disturbances ^24^.

### BOX 1: Quantification of the Net Biodiversity Effect on Stability

The net biodiversity effect on stability (NBES) can be quantified as the difference between observed and expected stability in experimental communities (Fig. 1). While observed stability is determined based on community responses to disturbance in the treatment relative to an undisturbed control, monospecific responses to disturbance in comparison to an equivalent undisturbed control are used to calculate an expected community response. This expected stability derives from the fundamental null assumption that each species shows proportionally the same response to the disturbance in monoculture and mixture, which we can sum according to species’ initial relative abundance in the assemblage.

In contrast to the net biodiversity effect on functioning, quantification of NBES relies on relative rather than absolute responses to an undisturbed control. It does not, therefore, allow the decomposition into selection and complementarity effects, which are based on absolute species responses^19^.

To calculate NBES, define for any mixture:

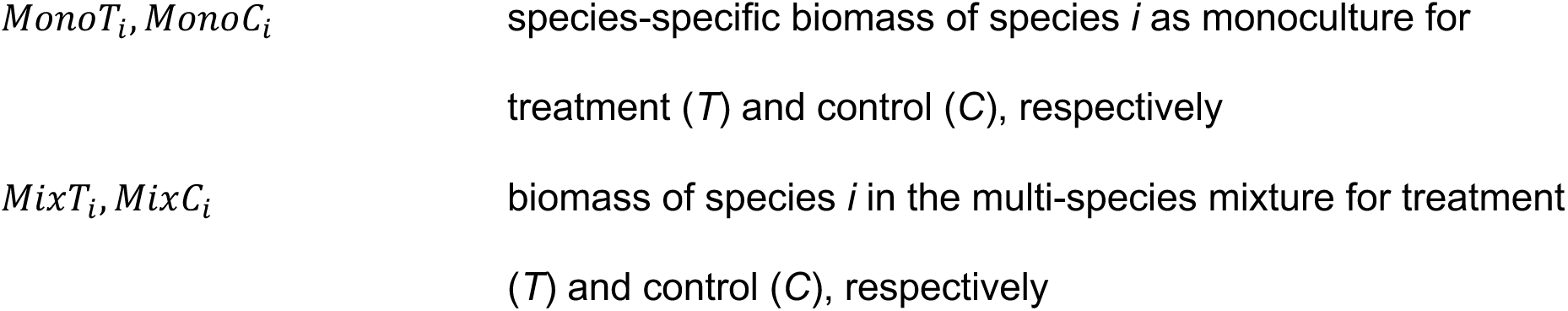
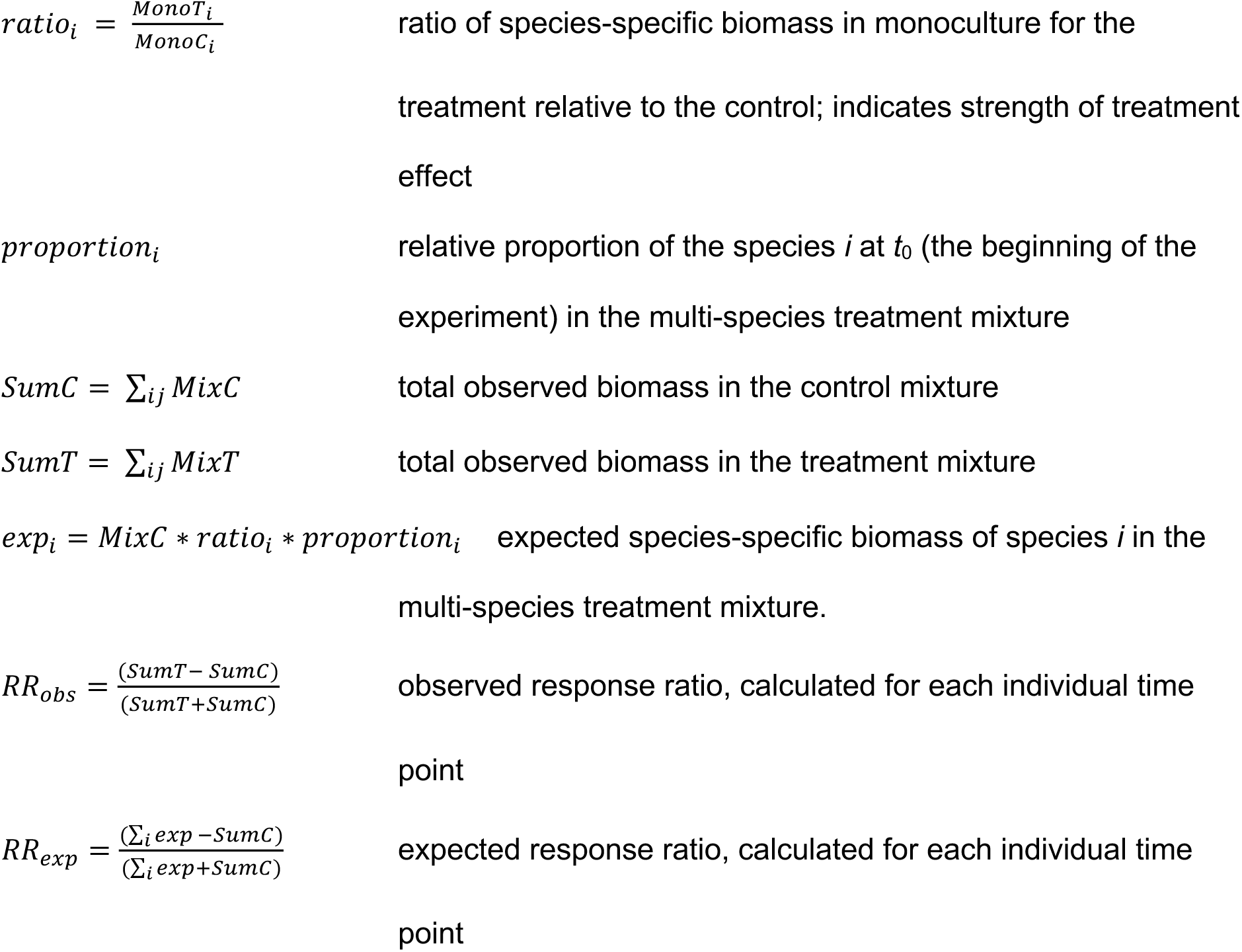

The NBES is then the difference between observed and expected stability (Fig. 1), estimated from the observed response ratio (*RR_obs_*), and expected response ratio (*RR_exp_*). In other words, the NBES is obtained from the difference between the observed and expected stability metric of interest. In this study, we assess the NBES primarily (though not exclusively) using the OEV index^20^. Calculation of the net biodiversity effect on stability is, however, not limited to any particular stability metric, and can be assessed for any metric that is based on a comparison of a treatment response to a control, such as resistance or temporal variability (see *Methods* for a detailed description). The calculation of stability should, however, be based on standardised response ratios (RRs) instead of log-response ratios (LRRs), to allow for the possibility that species become locally extinct, which would result in an undefined LRR.

While the calculation of the NBES is analogous to that of the net biodiversity effect on functioning^19^, the interpretation of NBES is sign-dependent. If the pressure at hand decreases biomass, NBES > 0 corresponds to a positive deviation from predictions and a stabilising biodiversity effect (Fig. 1), whereas NBES < 0 indicates a negative deviation from predictions and, thus, a destabilising effect of biodiversity.

**Fig. 1.**
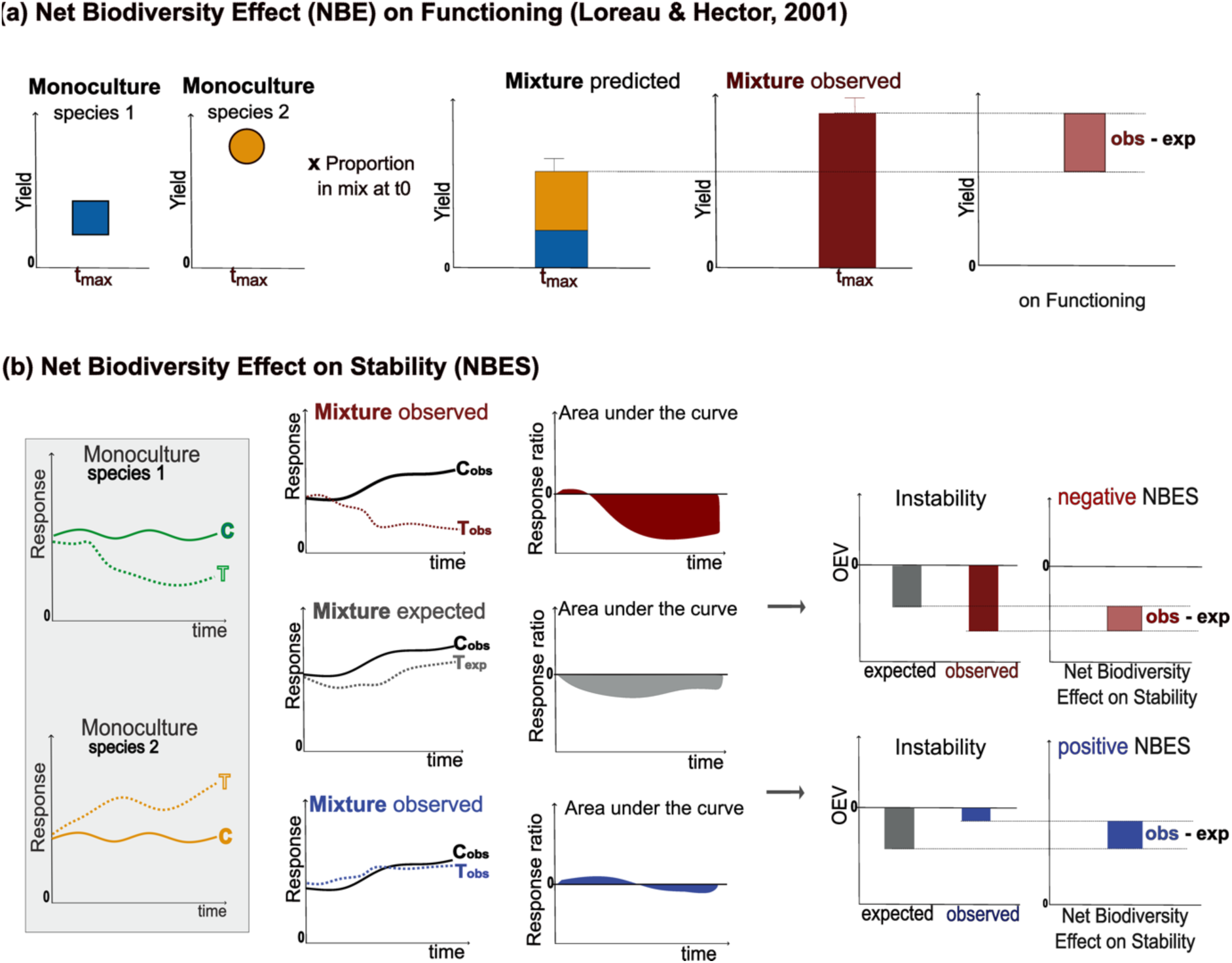
Quantification of the net biodiversity effect on functioning and stability. The net biodiversity effect (NBE) on functioning is calculated as the difference between an observed and an expected biomass yield based on monospecific yields at the end of an experiment (a). The NBE on stability is calculated as the difference between observed and expected stability. Expected stability is determined from the sum of species-specific response ratios in monocultures and their observed contribution to biomass in the equivalent undisturbed control mixture. Observed stability is estimated from the realised biomass in treatment, abbreviated as *T*, relative to the undisturbed control, abbreviated as *C*. As an example, we have marked the area under the curve of the observed and expected response ratio [based on Overall Ecological Vulnerability (OEV) in this example], which represents an integrative measure of (in)stability. Here, we concentrate on a typical, growth-reducing disturbance and decipher the two cases of lower observed stability than expected and thus a negative NBES (red colour), and higher observed stability than expected, and thus a positive NBES (blue colour) (b).

## Results

### Model simulations

In our simulation experiment, we assessed the NBES using a temperature-dependent Lotka-Volterra species competition model^25^ with different levels of competition strength (that is, no competition, intermediate, and strong competition) and different distributions of species-specific thermal optima (all species equal, each species different). The NBES was then assessed for every possible community combination of 2-5 species for three temperature disturbance treatments (pressure, fluctuations, combination) that were analogous to the temperature disturbance treatments in our microcosm experiment (please see *Methods* for a detailed description of the model).

Observed stability was significantly different from expected stability in most simulated communities across richness levels. However, the direction of the NBES differed among disturbance types and competition strengths (Fig. 2; Supporting Information Fig. S1). The strength of competitive interactions modulated the magnitude of the NBES. In simulations that included strong competition, communities exhibited larger variation in NBES, amplifying either stabilising or destabilising biodiversity effects depending on the species combination in the model run. When species had the same temperature optimum, the NBES was consistently positive and increased with increasing species richness (Supporting Information Fig. S1). In contrast, in model runs where species varied in their temperature optima and with strong competitive interactions, the NBES was more variable and included neutral and negative effects, sometimes resulting in non-monotonic dynamics as richness increased (Fig. 2).

**Fig. 2:**
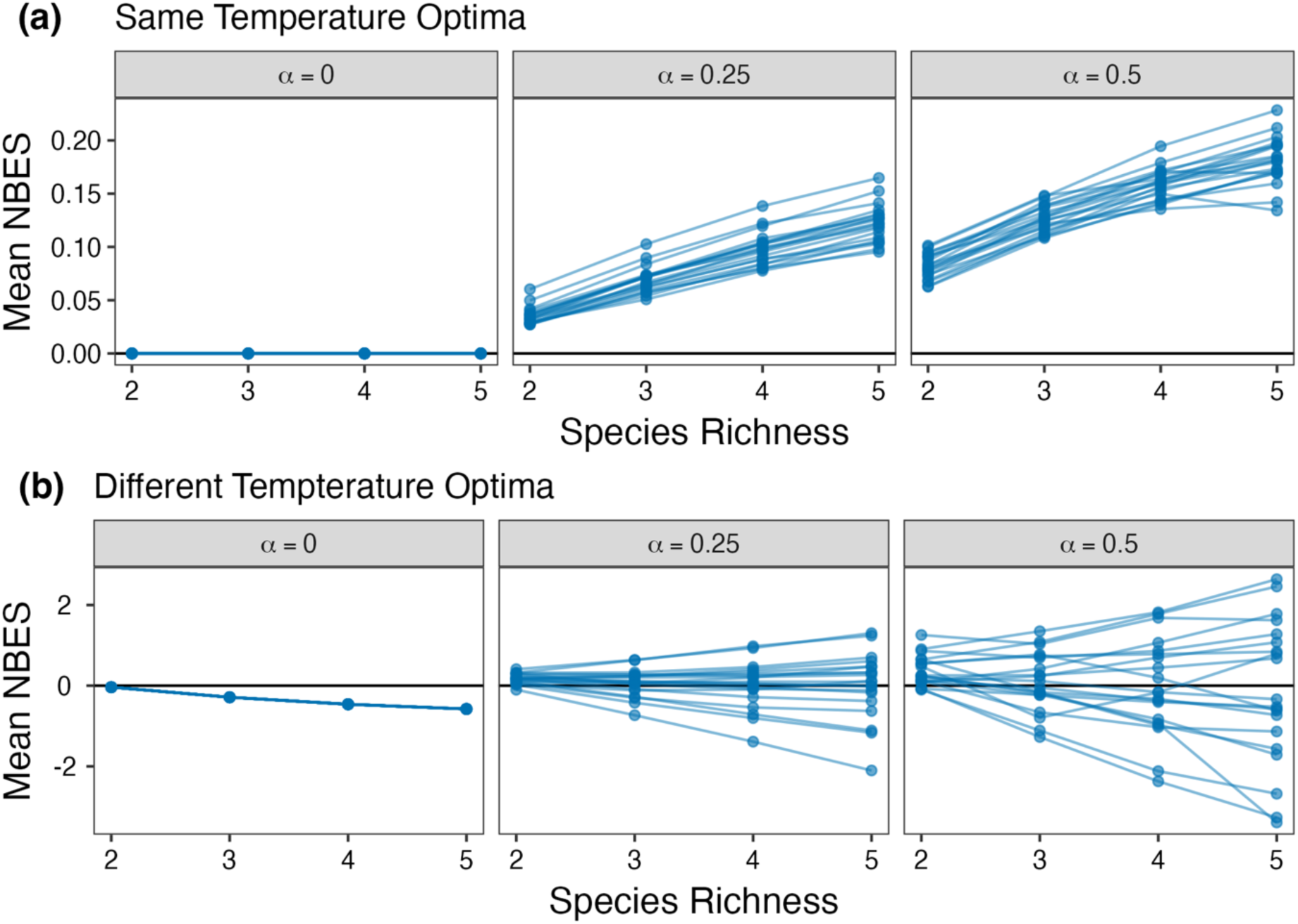
Results of model simulations for gradually increasing temperatures. Mean net biodiversity effect on stability (NBES) as a function of species richness for increasing temperature for model scenarios where (a) all species had the same temperature optimum (*b_opt_* = 17.5°C) and (b) all species had a different temperature optima (*b_opt_* ranging between 15 and 20 °C). Plots indicate increasing competition strength (α) from left to right. Here, we show only the results from the increasing temperature (press) treatment as the results for all three of our disturbance regimes were similar (for a complete overview of NBES in different disturbance regimes see Supporting Information Fig. S2).

Across competition strength increases, species in species-poor communities contributed more strongly to NBES than those in species-rich assemblages (Fig. 3), while their temperature preferences (*b_opt_*) determined the direction of the effect. Specifically, species with temperature optima at the extremes of the simulated range showed the greatest influence on NBES both positively and negatively. Under increasing temperatures, species with lower or slightly higher temperature optima showed a positive influence on NBES, whereas species with the highest temperature optima showed a negative influence on NBES. Under fluctuating temperatures, species with lower thermal optima showed both positive and negative influence on NBES depending on the competition strengths and community composition, whereas species with highest thermal optima showed a positive influence. The combined treatment mirrored the effects found for fluctuating temperatures but with greater variation in the influence on NBES for strong competition. In general, stronger competition resulted in more variable influences on NBES, reinforcing the role of interactions in shaping stability outcomes. Overall, these results highlight that biodiversity effects on stability are not uniform but are mediated by species interactions and temperature-dependent species performance.

**Fig. 3:**
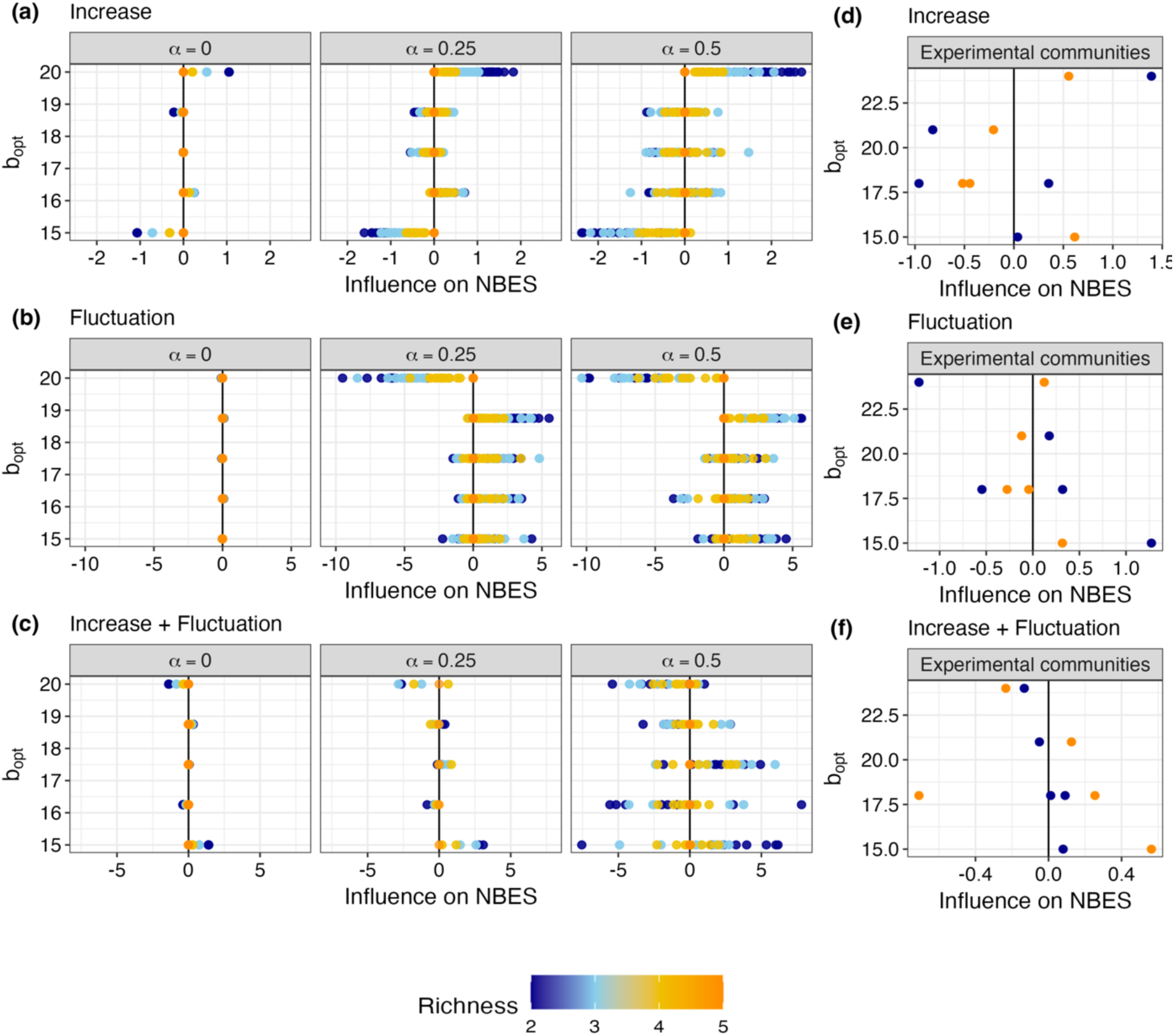
Relationship between the influence of species on the NBES and their temperature optimum *b_opt_* for model communities (a-c) and experimental communities (d-f). (a, d) Increasing temperature (b, e), fluctuating temperatures, and (c, f) increasing and fluctuating temperatures. All species had different temperature optima (*b_opt_* ranging between 15 and 20°C). Species temperature preferences (*b_opt_*) determined the direction of the NBES. Species richness is indicated by the color gradient.

### Microcosm experiment

In our microcosm experiment, we used five diatom species isolated from the North Sea and manipulated all possible species combinations across a gradient of richness (one, two, four, and five species). Cultures were exposed to one of four temperature treatments comprising a gradual temperature increase of 0.2°C per day, diurnal temperature fluctuations of ±3°C, the combination of gradual increase and fluctuations, and a constant temperature control, all starting at 17°C or with a mean of 17°C (see *Methods*).

Observed stability showed considerable variation across treatments and richness levels as well as associated species combinations (Fig. 4). Specifically, the majority of multi-species combinations showed positive or near-neutral deviations from the constant temperature controls, but four-species combinations deviated both positively and negatively from their controls. Most species monocultures showed negative deviations for disturbances involving fluctuations, whereas gradual temperature increases even led to positive deviations in some monocultures (Fig. 4).

**Fig. 4.**
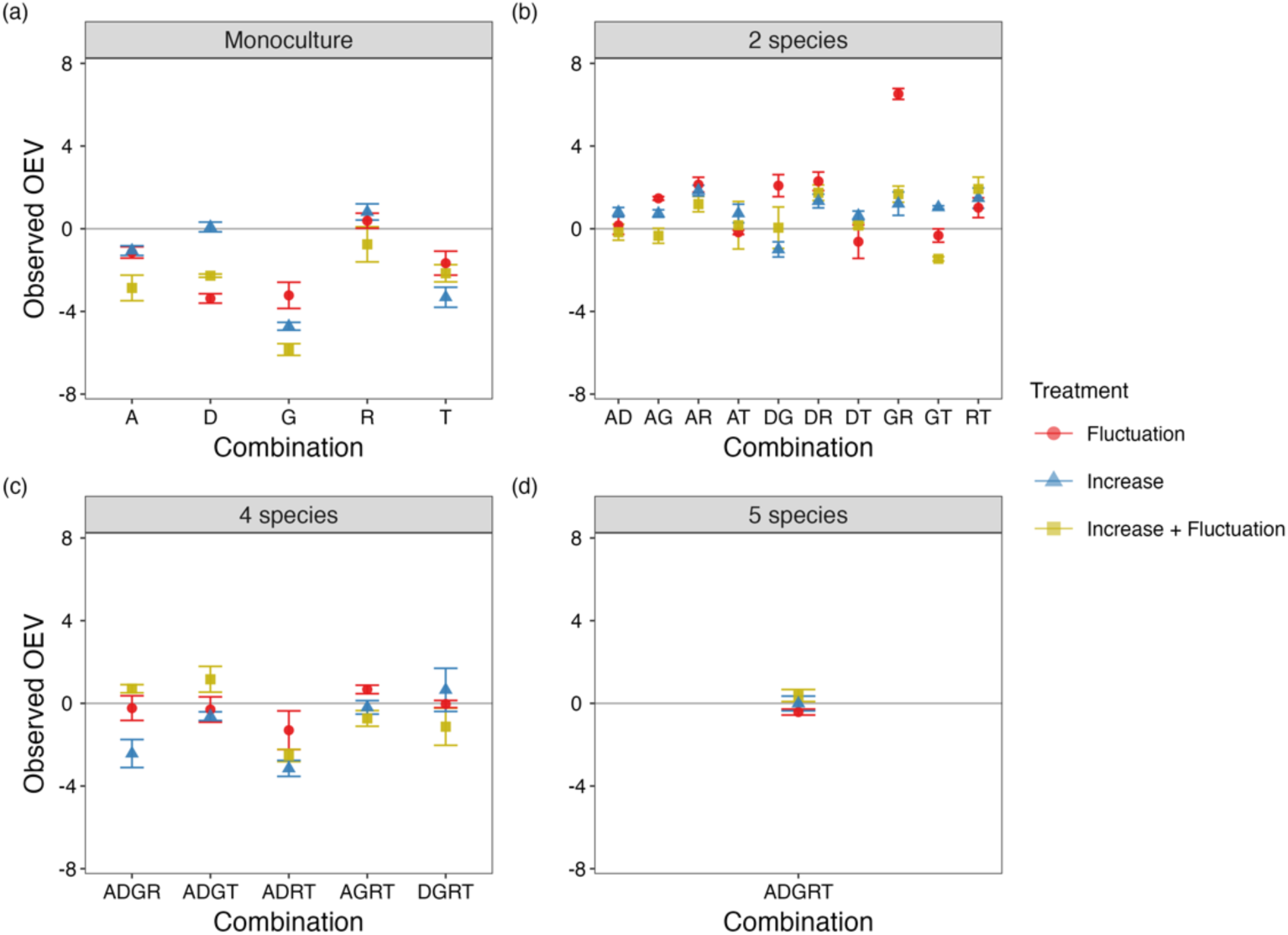
Observed instability, measured as Overall Ecological Vulnerability (OEV) for different species combinations. Mean (*n* = 3) observed instability ± SE for (a) monocultures, (b) two-species assemblages, (c) four-species assemblages, and (d) five-species assemblages. Higher absolute values indicate greater instability with positive and negative deviations. Different colours and shapes indicate different temperature treatments: diurnal fluctuations ±3 °C with a mean of 17 °C (red circles), temperature increase from 17 to 23 °C (blue triangles), and combined diurnal temperature fluctuation of ± 3 °C around an increasing mean (green squares). Phytoplankton species are abbreviated as A – *Asterionellopsis*, D – *Ditylum*, G – *Guinardia*, R – *Rhizosolenia*, T – *Thalassionema*.

Observed stability differed from expected stability in most communities of two, four or five phytoplankton species (Fig. 5a). The NBES was overall positive and differed significantly from zero (*t*-test across all compositions; *n* = 144, *t* = 7.1, *p* < 0.01). Moreover, both temperature treatments and species combinations, as well as their interaction, had a significant effect on NBES (Supporting Information Table S1, ANOVA, *p* < 0.05; Fig. 5b).

**Fig. 5.**
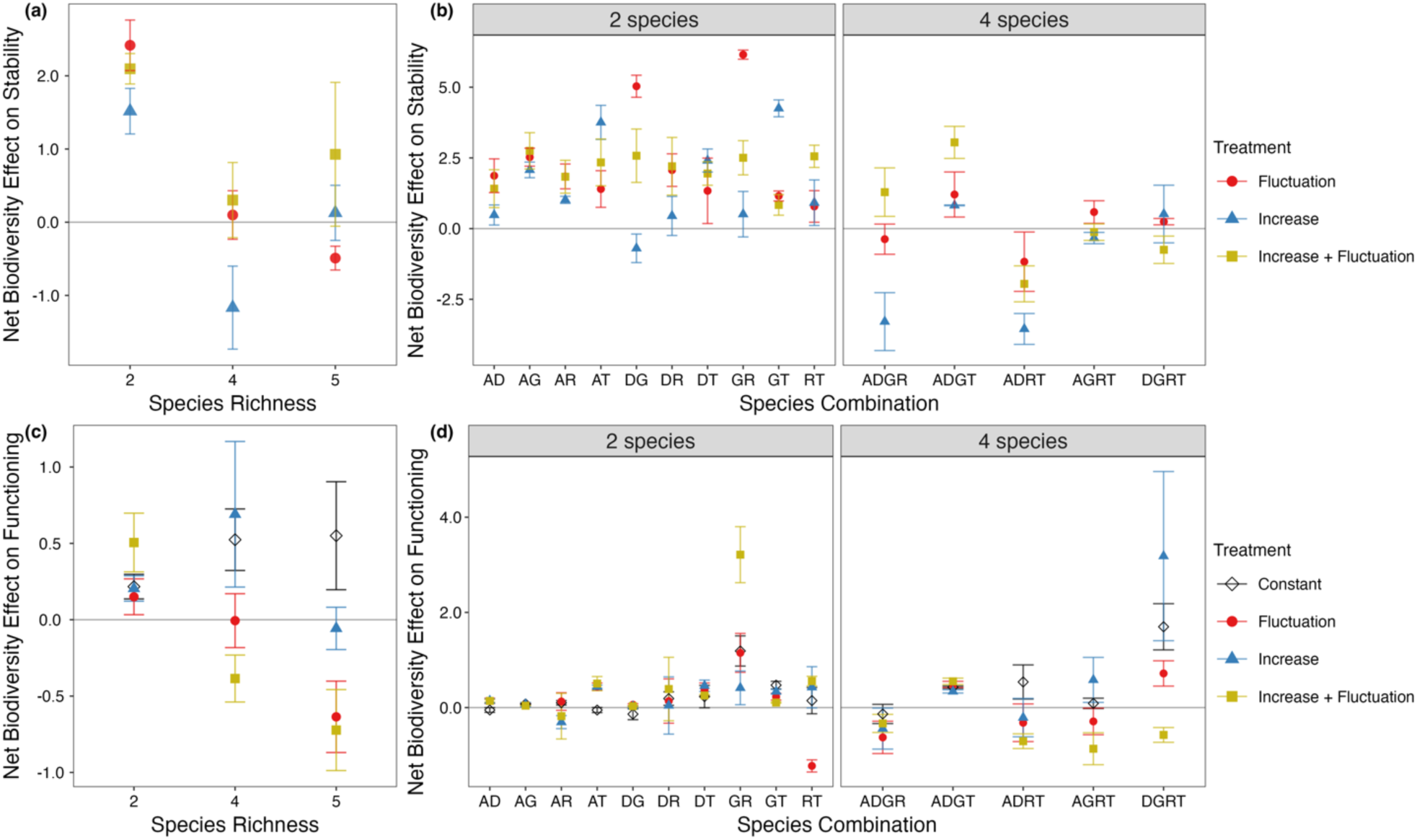
The net biodiversity effect on stability (NBES) for the different (a) species richness levels and (b) species combinations, and the net biodiversity effect on ecosystem functioning for different (c) species richness levels and (d) species combinations. Positive values of the net biodiversity effect on functioning and stability indicate a positive effect of biodiversity, and negative values a negative effect. Different colours and shapes indicate different temperature treatments. Phytoplankton species are abbreviated as A – *Asterionellopsis*, D – *Ditylum*, G – *Guinardia*, R – *Rhizosolenia*, T – *Thalassionema*. Each point indicates mean ± SE with *n* = 3 for plots (b), (d) and varying replicates for (a) and (c), where *n* = 30 for two species combinations, *n* = 15 for four-species combinations, and *n* = 3 for five-species combinations.

Specifically, for the combined temperature increase and fluctuations treatment the NBES was positive or neutral overall and decreased for four- and five-species assemblages. For both the increasing temperature regime and diurnal fluctuation temperature treatments, the NBES even turned negative at high richness levels (Fig. 5a). NBES of both four- and five-species assemblages differed significantly from those of two-species combinations (planned pairwise-comparisons; *p* < 0.05), but not from each other. Disentangling species effects on NBES revealed large differences among disturbance regimes. That is, species influence on the NBES aligned with their temperature optima (Fig. 3). Under gradually increasing temperatures, species with slightly higher temperature optima (between 18 and 21 °C) positively impacted NBES, while a gradual increase in temperature led to suboptimal conditions for species with extreme temperature optima, thereby negatively affecting NBES. Under fluctuating disturbances, species with higher temperature optima contributed mostly positively, while a lower temperature optimum had a negative influence on NBES. Overall, species in two-species communities exerted a stronger influence on NBES compared to those in multi-species assemblages of four or five species.

The net biodiversity effect on functioning varied significantly among species combinations, temperature treatments, and the interaction between species combinations and temperature treatments (ANOVA, *p* < 0.01, Supporting Information Table S1).

Specifically, the net biodiversity effect on functioning was always positive in the constant temperature control and increased with increasing species richness (Fig. 5c). In contrast, disturbance treatments decreased the net biodiversity effect on functioning as species richness increased, and even turned it negative in most cases for four- and five-species combinations (Fig. 5d). One exception was the gradual temperature increase with a positive net biodiversity effect on functioning for four-species assemblages due to an increase in total biomass at the end of the experiment driven by *Thalassionema* (Supporting Information Figs. S2-4). Overall, the net effects of biodiversity on stability and functioning correlated positively (Spearman-Rank correlation, *R* = 0.35, *p* < 0.01).

## Discussion

Community-wide functional stability could not be predicted by species-specific responses to the same environmental change in monocultures. Rather, simulated communities showed both lower and greater stability than expected, whereas experimental communities showed overall greater stability than expected (that is, an overall positive NBES; partly accepting H1). NBES varied widely in direction and magnitude in the simulations, while in the experiment NBES was highest for two-species combinations and decreased at higher species richness levels (partially accepting H2). NBES varied among species combinations and temperature treatments according to species thermal optima in both simulations and the experiment, and even became negative for some (accepting H3). Finally, the magnitude of the NBES varied with competition strength in the simulations (accepting H4). Our approach enabled us to quantify the amount of additional stability derived from biodiversity, in terms of both richness and composition, therefore providing novel and important insights into the biodiversity – stability relationship.

We first tested the NBES approach using simulated data with varying competition strengths, disturbance regimes, and distributions of thermal optima. Our framework enabled consistent estimation of NBES across all modelled conditions, including cases with minimal or neutral biodiversity effects. The direction and magnitude of the NBES depended on competition strength and species’ temperature optima. When species shared a favourable optimum, NBES was consistently positive and increased with increasing competition strength, whereas greater variation in thermal preferences led to more variable NBES outcomes with much greater magnitude. This suggests that the effect of biodiversity on stability depends on how well the traits of a community’s component species match the prevailing environmental conditions ^26,27^. That is, how well the temperature optima match the regime temperature. Strong competition and high diversity in species temperature optima sometimes led to non-monotonic dynamics (Fig.3 & Supporting Information Fig. S1). This was more often the case for our temperature increase treatment than for temperature fluctuations and temperature increase and fluctuations in combination. Here, species with stronger competitiveness showed greater variation in their influence on the NBES, as interactions tend to modulate species responses to disturbance^24^. Moreover, we find greater variation in the direction of the NBES with increasing species richness, suggesting that more diverse communities may exhibit a wider range of stability outcomes. This is consistent with the theoretical work of Ives and Carpenter (2007), who showed that species-rich systems are more likely to show a broader range of diversity-stability relationships in the face of disturbance because interspecific interactions can both amplify and dampen stability effects.

In our microcosm experiment, the NBES was overall positive, which we expected based on previous reports that biodiversity dampens the variability of emergent properties of the community in fluctuating environments ^5,6,9,28^. Contrary to expectations that the NBES would increase with increasing species richness, however, we found that the mean NBES decreased for higher levels of species richness (Fig. 5). One reason for this may be the lower resistance of species-rich assemblages to disturbance treatments than expected from monoculture and compared to two-species assemblages, resulting in a negative trend for the NBES for resistance along the species richness gradient (Supporting Information Fig. S3). If species-rich systems have a greater chance of harbouring species that grow well under undisturbed conditions, as suggested by the biodiversity insurance hypothesis^13^, they also have a greater chance of losing this growth potential under disturbance. This is especially true if there exists a trade-off between species growth and resistance to disturbances^10^. This investment in growth rather than resistance in species-rich communities may allow for compensatory fluctuations due to asynchronous responses among species over time^6,29,30^ despite the sustained disturbance pressure. Consistent with this, we also found lower NBES for temporal variability (that is higher temporal stability) in species-rich assemblages than in two-species assemblages (Supporting Information Fig. S3), indicating that observed variability was lower than expected in species-rich assemblages. This suggests a negative correlation between resistance and temporal stability (that is, low resistance and high temporal stability with increasing species richness). Consistent with this, Pennekamp *et al.*^8^ also reported negatively covarying stability metrics (that is, high temporal stability and low resistance), which resulted in a ‘U’-shaped relationship between overall ecosystem stability and species richness under gradual warming. Combined with our results, this highlights the necessity of considering the multidimensional nature of stability^2,31^.

The net biodiversity effect on functioning largely mirrored those for stability in our microcosms, decreasing with increasing species richness when exposed to disturbance treatments, but increasing in the constant temperature controls of our experiment. The positive relationship between productivity and species richness, as in our controls, is a frequent observation in diversity manipulations and often attributed to the fact, that in more diverse ecosystems different species use resources in different ways, leading to more efficient resource utilisation^10,13,32^. Most studies in the context of biodiversity and ecosystem functioning have been conducted in undisturbed systems^32,33^, which may not fully capture the dynamics under varying environmental conditions. However, disturbances have the potential to disrupt this biodiversity–productivity relationship, and even turn it negative, due to the trade-off between resistance to disturbance and growth rate^10^. This context-dependency of the net biodiversity effect on functioning in our experiment suggests a positive correlation between productivity and stability in our experimental system.

Overall, both simulations and experiments revealed high heterogeneity in the NBES within richness levels and among species combinations, reflecting diverse community responses to disturbance regimes (Figs. 2, 6). Much of this variation was driven by species’ thermal optima—species whose traits matched the prevailing temperature regime tended to have a more positive effect on the NBES. This variation in the influence of different species on the NBES based on their thermal optima suggests a strong idiosyncratic compositional effect. That is, rather than having a uniform response across the entire community, individual species may exhibit distinct responses to the different temperature regimes and, therefore, affect stability in different ways in different contexts^34–37^. Such variation aligns with known differences in species’ thermal responses and metabolic rates^21,22,38^. For example, gradual warming often reduces growth^23^, as shown in *Asterionellopsis, Guinardia*, and *Thalassionema* monocultures, while *Ditylum* and *Rhizosolenia* performed better (Supporting Information Fig. S2). Diurnal fluctuations led to lower biomass in most species, consistent with predictions from non-linear averaging^39^. While temperature fluctuations around a gradually increasing mean temperature represented a brief exposure to temperatures close to the thermal optimum for many species^40^, this resulted in higher biomass production than under ‘simple’ diurnal fluctuations, but negative deviations from constant temperature controls for most species (and thus a negative influence on NBES). These differences in species responses may also reflect different adaptation mechanisms of species, such as reductions in cell size and increased nutrient storage^35,41^, or temporal differences in these adaptation mechanisms^42,43^.

Disturbance regimes might also affect the outcome of interspecific interactions differently. Rising temperatures may lead to increased diversification of species interactions in communities because of buffering mechanisms to environmental change^29,36^. Specifically, rising temperatures may enhance competition between species due to the critical role of metabolic traits in determining the impact of temperature changes on interspecific interactions^38^. Such increased competition may have manifested in a negative NBES for higher levels of species richness in our experiment, which is consistent with results from the model simulations for strong competitive interactions (Fig. 3). In contrast, temperature fluctuations may increase the likelihood of long-term species asynchrony, thereby facilitating community and ecosystem stability, despite potential short-term destabilising effects^13,29,30^. This is consistent with our results of the experiment and model simulations for disturbances involving fluctuations, where we see greater variation in species contributions to NBES based on their thermal optima.

We applied the NBES approach to simulated multi-species communities and experimentally to microcosm communities with undisturbed controls. Our approach could, however, also be applied to observational data if post-disturbance dynamics of both mono-specific and species-specific responses in the community are available. Specifically, if observational time series data include both monoculture dynamics and species-level contributions within mixed communities, NBES can be calculated by comparing observed community stability against expectations derived from single-species responses, enabling application of the framework across different ecosystems and disturbance scenarios.

Given the multitude of experimental contexts and different outcomes in combination with model simulations using Lotka-Volterra dynamics, our experimental design was ideal for testing the difference between expected and observed stability. Our simulations cover a range of competition strengths and distribution of thermal optima, showing that the NBES can be both positive and negative, depending on community and trait composition. However, our microcosm experiment was based on a single system using highly productive marine primary producers. We therefore encourage further studies to determine the NBES for other organism groups in other systems to explore potential trade-offs between both stability and productivity^10,44,45^, and the importance of interaction type (competition vs. mutualism).

The various effects of different disturbance treatments on the net effect of biodiversity on stability and functioning highlight the need to consider a wide range of disturbance types and characteristics when studying the consequences of biodiversity. Given the scale-dependency of biodiversity^46^, it is essential to broaden the spatial and temporal scope of biodiversity research beyond individual systems (*e.g.*, food webs) and specific time frames. In particular, the introduction of a greater number of species and species richness levels could further elucidate the dynamics of multi-species assemblages. This will facilitate a deeper understanding of the underlying mechanisms that govern the biodiversity-stability relationship and how they may change over space and time.

## Conclusion

By quantifying the additional stability provided by biodiversity under three temperature change scenarios using both model simulations and a microcosm experiment, we show that stability emerges as a community property that is influenced not only by species richness but also by species identity. Different species played key roles in determining community stability under different disturbance regimes depending on their thermal preferences. This suggests that species loss may have unforeseen and severe consequences for ecosystems under climate change and highlights the context-dependency of biodiversity effects on stability. In practical terms, our findings highlight the importance of maintaining biodiversity to enhance ecosystem resilience to external forcing and maintain productivity, and thus provide crucial insights for managing ecosystems in times of biodiversity crisis and changing environmental conditions.

## Methods

### Model simulations

We simulated a five-species community using a continuous Lotka-Volterra model with temperature-dependent vital rates^25^. We used a continuous temperature-dependent Lotka– Volterra model (instead of a time-discrete model^47^) to capture the time-varying effects of temperature on population dynamics. In the model, the change of biomass for each species is described as

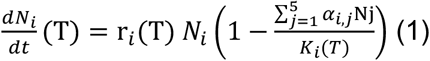

where *r_i_(T)* corresponds to the species-specific biomass, *r_i_(T)* is the species-specific temperature-dependent intrinsic growth rate, which is given by the difference between the birth rate (*b*_0_,*i*(*T*)) and death rate (*d*_0,i_(T)), *K_i_*(*T*) the species-specific temperature-dependent carrying capacity, *α_i,j_* the competition strength between species *i* and species *j*, and *N_j_* the biomass of species *j*.

The carrying capacity *K_i_(T)* of species *i* is given as

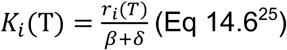

where *β* and *δ* are the density dependent constants that were set to 0.025, respectively. Temperature dependence (*T*) was incorporated into the birth and death rate:

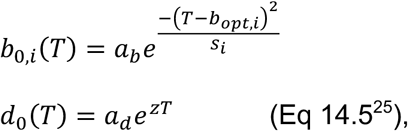

where *a_b_* and *a_d_* are the intercepts of birth and death rates, respectively, *b_opt,t_* is the species-specific temperature at which the intrinsic growth rate is highest, *s* scales the width of the temperature response of birth rate, *z* is the slope of the death rate and scales the effect of temperature (in °C) to mimic the Arrhenius relationship ^25^. Parameter values were so that the community would reach equilibrium biomass within 150-time steps (see Supporting Information Table S2 for an overview of introduced parameter values).

The interspecific competition terms *α_i,j_* were drawn from a one-sided normal distribution *α* > 0, so that all interactions were competitive with asymmetric interactions between any two species in the model. A higher value for α indicates a stronger effect by species *i* on species *j*, whereas a low value of *α* indicates a small effect of species *i* on species *j*. We varied interspecific competition terms by introducing differing standard deviations (SD) of competition values *α_i,j_* while the mean remained constant (*µ* = 0).

Specifically, we tested three competition strengths in the communities:

i. no competition(SD = 0),
ii. intermediate competition strength (SD = 0.25), and
iii. high competition strength (SD = 0.5).

The distribution of species’ temperature optima (*b_opt,i_* within communities was evenly distributed along specific temperature gradients to represent varying levels of diversity in species responses. We specifically simulated the following two scenarios:

i. All species had the same *b_opt,i_* (17.5°C), simulating a homogenous response.
ii. All species had different *b_opt,i_*, that were evenly distributed between 15°C and 20°C.

We simulated species within each community as monocultures and in every possible combination of 2 – 5 species. Each of the species assemblages was then exposed to a disturbance and a control run for 150 time steps. The disturbance regimes comprised of a temperature increase from 15 – 20°C (press), temperature fluctuations from 15 – 20°C within one time step with a mean temperature of 17.5°C (fluctuations), a combination of temperature increase from 15 – 20°C with fluctuations of ± 2.5 °C, and an undisturbed control at 17.5°C constant. The temperature scenarios commenced at the first time step to be consistent with our microcosm experiment (see below). To account for the stochasticity when drawing the competition values *α_i,j_*, each combination of disturbance type (press, fluctuations, combination), competition strength, and temperature optima distribution (*b_opt,i_*) was repeated 20 times, resulting in a total of 480 scenarios (see Supporting Information Fig. S6 for an exemplary model run).

### Microcosm experiment

To quantify the NBES empirically in real communities, we conducted a microcosm experiment on marine phytoplankton with different levels of species richness, from monocultures to multi-species assemblages of five species, and crossed that gradient with four temperature treatments (Fig. 6). Phytoplankton communities are an ideal system to study the effects of biodiversity on stability to disturbance because of their short generation times and high sensitivity to environmental changes ^3,48,49^, while supporting high species diversity ^50^. We used five different phytoplankton species isolated from the North Sea in summer 2017 that differed considerably in cell volume: *Asterionellopsis glacialis* (252 µm^3^),

**Fig. 6.**
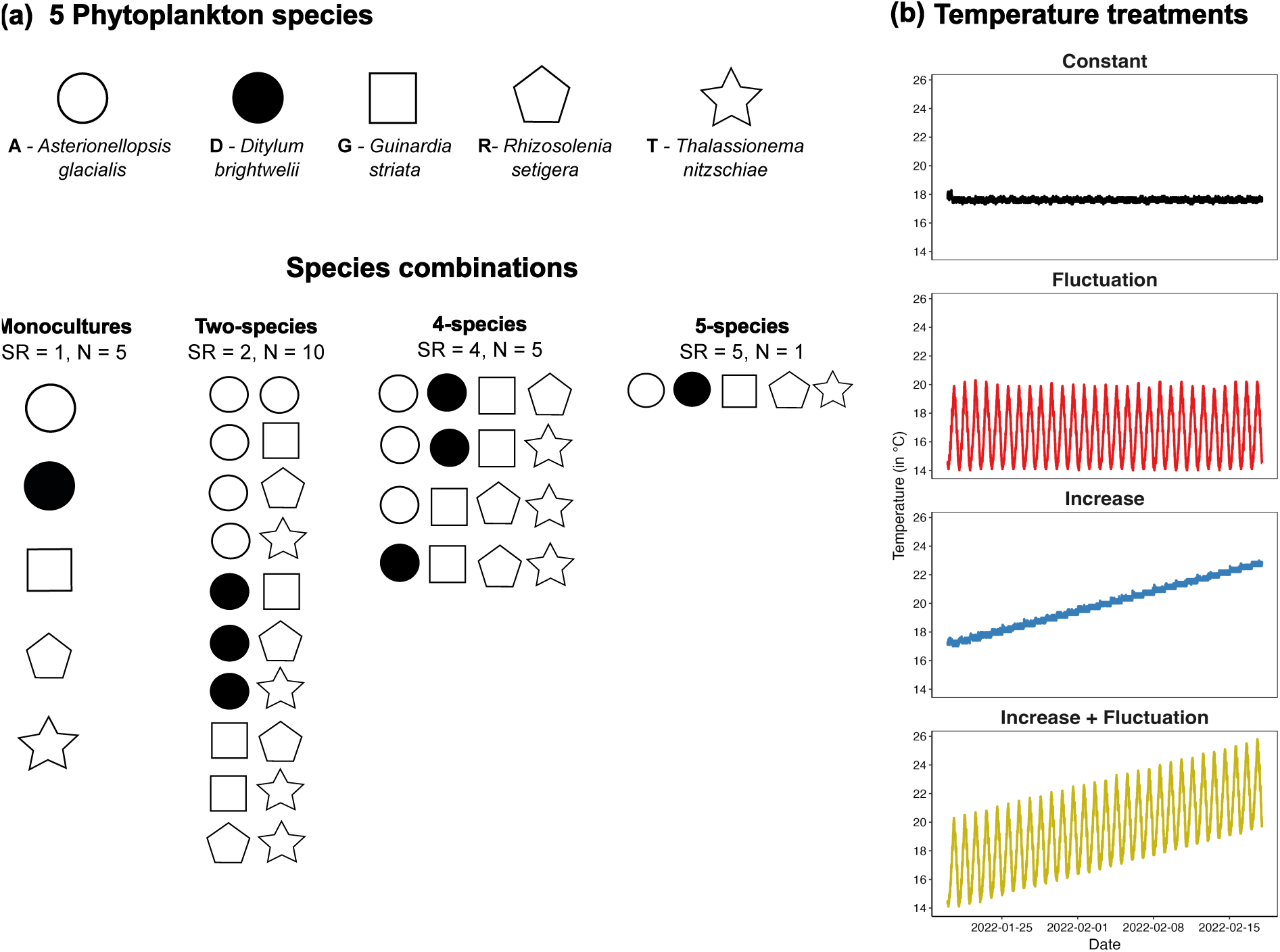
Overview of our experimental design. Our microcosm experiment comprised (a) four levels of species richness, comprising monocultures of each respective species, two species in combination, four species in combination and all five species in combination crossed with (b) four temperature treatments, with each temperature treatment-species combination replicated three times. Each colour represents one species. Displayed temperature curves are measured temperatures for the different treatments over the experimental period for one replicate mesocosm each. All temperature treatments started at 17°C or with a mean temperature of 17°C and consisted of a constant temperature treatment with 17°C, a diurnal temperature fluctuation ± 3°C from 14 to 21°C, a fluctuation combined with increasing temperature where the temperature fluctuated ± 3°C around the increasing mean, and a temperature increase from 17 to 23°C.

*Ditylum brightwelii* (2204 µm^3^), *Guinardia striata* (5667 µm^3^), *Thalassionema nitzschioides* (7236 µm^3^), and *Rhizosolenia setigera* (205215 µm^3^). Mean cell volume was estimated from 20 individuals of each diatom species at the beginning of the experiment under an inverted microscope. We chose these species because of their large variation in cell size, which is known to be a master trait in phytoplankton, influencing differences in growth rates, responses to temperature, and nutrient prevalence ^51–53^. In addition, diatoms play a key role in marine primary production worldwide^54^.

Prior to the start of the experiment, cultures were grown at 12 °C in F2 medium at 25 PSU at ∼200 µmol s^-1^ in a 12:12 dark: night cycle. To create disturbance regimes with negative effects on species growth rates, each temperature treatment started at 17°C or with a mean temperature of 17°C and reached maximum temperatures of 23°C, thereby exceeding (or coming close to) the average temperature optimum of marine phytoplankton species from the North Sea at 18°C ^40^. In our experiment, species temperature optima ranged from 15-24°C with 18 °C for *Asterionellopsis*, 21 °C for *Ditylum*, 15°C for *Guinardia*, 18°C for *Rhizosolenia*, and 24°C for *Thalassionema* (assessed as maximum biomass along a temperature gradient). Specifically, we introduced four different temperature treatments (Fig. 6): a constant temperature control at 17 °C, a diurnal fluctuation treatment from 14 – 20°C in a 12:12 cycle, a warming treatment with temperatures increasing by 0.2 °C per day from 17 °C to 23 °C, and diurnal temperature fluctuations combined with warming with a fluctuation of ± 3°C around an increasing mean of 0.2 °C per day. We chose a gradual increase in temperature and temperature fluctuations as disturbance treatments on the one hand because pulse disturbances have received considerably more attention than other forms of disturbance in stability research^11,55^. On the other hand, diurnal temperature fluctuations and fluctuations around an increasing mean represent more realistic disturbances compared to pure press or pulse disturbances ^31^ and compared with other fluctuation frequencies^56^.

Our experiment comprised four levels of species richness (Fig. 6), each of which was replicated three times. Species richness levels comprised monocultures of each of our five focal species—two species in combination, four species in combination, and all five species in combination. We estimated the density of our pre-cultures photometrically and added a fixed fraction to the medium to initiate all mono- and multi-species cultures at an optical density of ∼0.05. Monocultures and multi-species communities (containing two to five species) were assembled using a substitution design. All cultures were grown in a total of 35 ml medium and in sterile 50 ml polyethylene cell culture flasks (Sarstedt).

The experiment was conducted in the ‘Planktotrons’ indoor mesocosm facility ^57^, custom-tailored for plankton experiments at the Institute for Chemistry and Biology of the Marine Environment (ICBM). Temperature regimes were created using water baths, thereby controlling the temperatures with the heating and cooling units of the Planktotrons (Fig. 6). The experiment lasted 30 days and all treatments started at day 0 on 19 January 2022. The samples were kept in a 12:12 light: dark cycle at ∼150 µmol s^-1^. Nutrient concentrations were chosen to be close to ambient conditions (that is, measured dissolved nutrient concentrations were N 18 µmol, Si 17 µmol, P 1.5 µmol on day 0).

For sampling, experimental units were taken out of the water bath, stirred gently to homogenise the culture and a sample of 0.5 ml transferred into 48-well plates (Sarstedt). We sampled each experimental unit every sixth day and fixed samples with 50 µl of 10% Lugol’s iodine, resulting in a total of six time points over 30 days (that is, days 0, 6, 12, 18, 24, 30).

We derived the biomass of individual species by measuring their density under an inverted microscope at 200 – 400x magnification and estimating their biovolume ^58,59^.

### Stability calculation

We quantified the net biodiversity effect on stability (NBES; Box 1) based primarily on the integrative metric of Overall Ecological Vulnerability (OEV^20^). The OEV is calculated as the area under the curve of the total biomass in disturbed treatments relative to the undisturbed control. Here, we allow negative and positive values of the area under the curve, thus allowing both negative and positive deviations from controls to cancel ^47^. Instead of log-response ratios (LRRs), the calculation of OEV is based on standardised response ratios (RRs) to allow for the possibility of local extinctions of species. We then calculated NBES as the difference between observed OEV and expected OEV (Box 1), assessed as the area under the curve of the expected and observed standardised response ratios (RRs), respectively, that is, as:

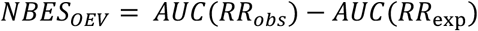

A positive NBES then indicates a stabilising effect of biodiversity, as measured instability is lower (less negative) than expected (that is, observed OEV – expected OEV is positive).

For data from the microcosm experiment, as well as using OEV as the basis of our assessments, we assessed the NBES for both resistance and temporal variability. The NBES for resistance was assessed as the difference between observed and expected RR at the first sampling (*t*_1_) after the commencement of disturbance, that is on day 6:

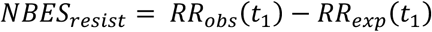

whereas the NBES for temporal variability was determined from the difference in the coefficient of variation (CV) from observed and expected *RR* over time:

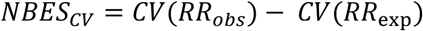

with 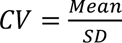

### Data analyses

All analyses were conducted using the R programming language in R version 4.4.3^60^.

For calculation of the area under the curve, we applied the auc() function within the MESS package ^61^ using linear splines and allowing for negative areas. For analysis we used packages available within the tidyverse ^62^, ggpubr ^63^, cowplot ^64^, and here ^65^. Conceptual figures were created using Inkscape 1.2 (version dc2aeda, 2022-05-15).

To test for a species identity effect of individual species on the NBES, we analysed the relationship between species temperature optima and the influence on NBES. Specifically, we calculated the species-specific influence on NBES as *X̄* − *Ā*, where *X̄* represents the grand mean over all species combinations per richness level and *Ā* is the species specific mean over all combinations that include species *A.* Species had a positive effect on the NBES when the NBES in combinations that included the specific species (*Ā*) was greater than the average NBES (*X̄*).

To test the effect of temperature treatments and species combinations on NBES in experimental communities, we conducted separate Analysis of Variance (ANOVA) tests, with species combinations and temperature treatment as independent variables. Using species combinations instead of species richness levels ensured a balanced analysis by introducing the same number of replicates. In case of a significant interaction effect of species combinations and temperature treatments, we conducted planned comparisons to compare species richness levels. Here, species richness levels were treated as factors to account for the differences in observation numbers.

The net biodiversity effect on functioning was calculated following Loreau & Hector (2001) from the observed and predicted biomass yield at the end of the experiment (*t*_max_). Similar to the NBES, we tested for an effect of temperature treatments and species combinations on the net biodiversity effect on functioning using ANOVA, with species combinations and temperature treatment as independent variables.

## Acknowledgements

C.K., H.H., M.S., and T.S. were supported by DFG funding (HI848/29-1). H.H. was supported by HIFMB, a collaboration between the Alfred-Wegener-Institute, Helmholtz-Centre for Polar and Marine Research, and the Carl-von-Ossietzky University Oldenburg. D.B. was supported by BMEL funding (2819HS015). We thank Heike Rickels and Christian Spindler for nutrient measurements and technical support. We thank Enja Dekena for helping with sample processing and Anna Lena Heinrichs for useful discussions.

