## Supporting Information for "Quantifying the net effect of biodiversity on stability"

**Supporting Information Table S1. The effect of species combinations and temperature treatments on Net Biodiversity Effects.** Results of the two-way ANOVA on the effect of species composition (Combination) and temperature treatments (Temp) on the net biodiversity effect (NBE) on Stability and the NBE on Functioning. Significant effects are indicated in bold.

| **Variable** | **NBE on Stability** | | | **NBE on Functioning** | | |
|---|---|---|---|---|---|---|
| | **DF** | **F-value** | **p-value** | **DF** | **F-value** | **p-value** |
| Temp | 2 | 11.24 | **<0.01** | 3 | 2.72 | **0.04** |
| Combination | 15 | 15.34 | **<0.01** | 15 | 8.65 | **<0.01** |
| Temp: Combination | 30 | 5.19 | **<0.01** | 45 | 3.13 | **<0.01** |

**Supporting Information Table S1. Overview of parameter values and variables used in the model simulations.**

| Variable or parameter | Description | Value/ Formula | Reference |
|---|---|---|---|
| **Species level variables and parameters** | | | |
| $\boldsymbol{N_i(t0)}$ | The biomass of species *i* at time *t0* | 0.1; same for all species in all simulated communities. | |
| $\boldsymbol{b_{opt,i}}$ | Temperature optimum of birth rate of species *i* | Either all species had the same $\boldsymbol{b_{opt,i}}$ = 17.5°C or all species had different $\boldsymbol{b_{opt,i}}$ that were evenly distributed between 15°C and 20°C. | |
| $\boldsymbol{a_{b,i}}$ | Birth rate at optimum temperature | 1; same for all species in all simulated communities. | |
| $\boldsymbol{s_i}$ | Width of temperature response curve of birth rate. | 30; same for all species in all simulated communities. | |
| $\boldsymbol{a_{d,i}}$ | Constant of temperature-death rate function | 0.01; same for all species in all simulated communities. | |
| $\boldsymbol{z}$ | Slope of temperature-death rate function. | 0.2; same for all species in all simulated communities. | |
| $\boldsymbol{\beta, \delta}$ | Density dependent constants for species carrying capacity. | 0.025; same for all species in all simulated communities. | |
| **Community level parameters** | | | |
| $\boldsymbol{\alpha_{ij}}$ | The strength of interspecific interaction between species *i* and *j*. | Absolute value of draws from a normal distribution with mean 0 and standard deviation $\boldsymbol{\alpha_{ij}\_sd}$ | |
| $\boldsymbol{\alpha_{ij}\_sd}$ | Strength of interspecific competition in a community. | Sequence of 0, 0.25, 0.5. Low values create a community with weak competitive interactions, high values create a community with strong interactions. | |
| ***S*** | Species richness | 5, same for all communities | |
| ***T*** | Temperature | Control = 17.5°C<br>Increase = 15 (Temperature minimum) and 20°C (Temperature maximum)<br>Fluctuations = 17.5°C (Mean temperature) $\pm$ 2.5 °C | |

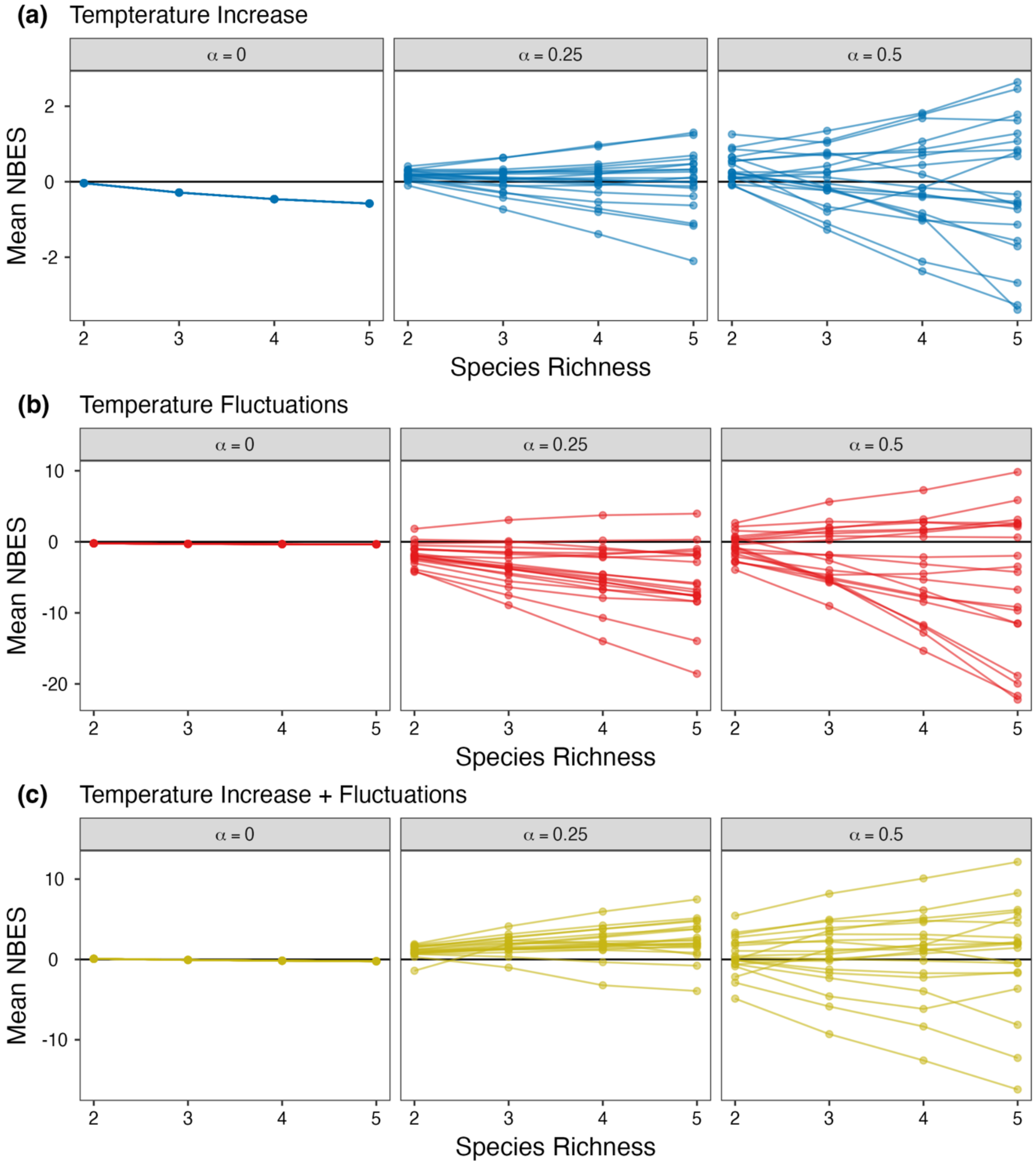


**Supporting Information Fig. S1**: Results of the model simulations for communities where species had different temperature optima (TOpt ranging between 15-20 °C). Mean net biodiversity effect on stability (NBES) for each model run and species richness level under (a) increasing temperatures (press), (b) temperature fluctuations, and (c) the combination of temperature fluctuations and temperature increase. The competition strength determined the magnitude of the NBES, while the direction of the NBES was dependent on the temperature optima distribution. Different colors indicate different disturbance regimes, different facets indicate different competition strength where higher values indicate stronger interspecific competition.

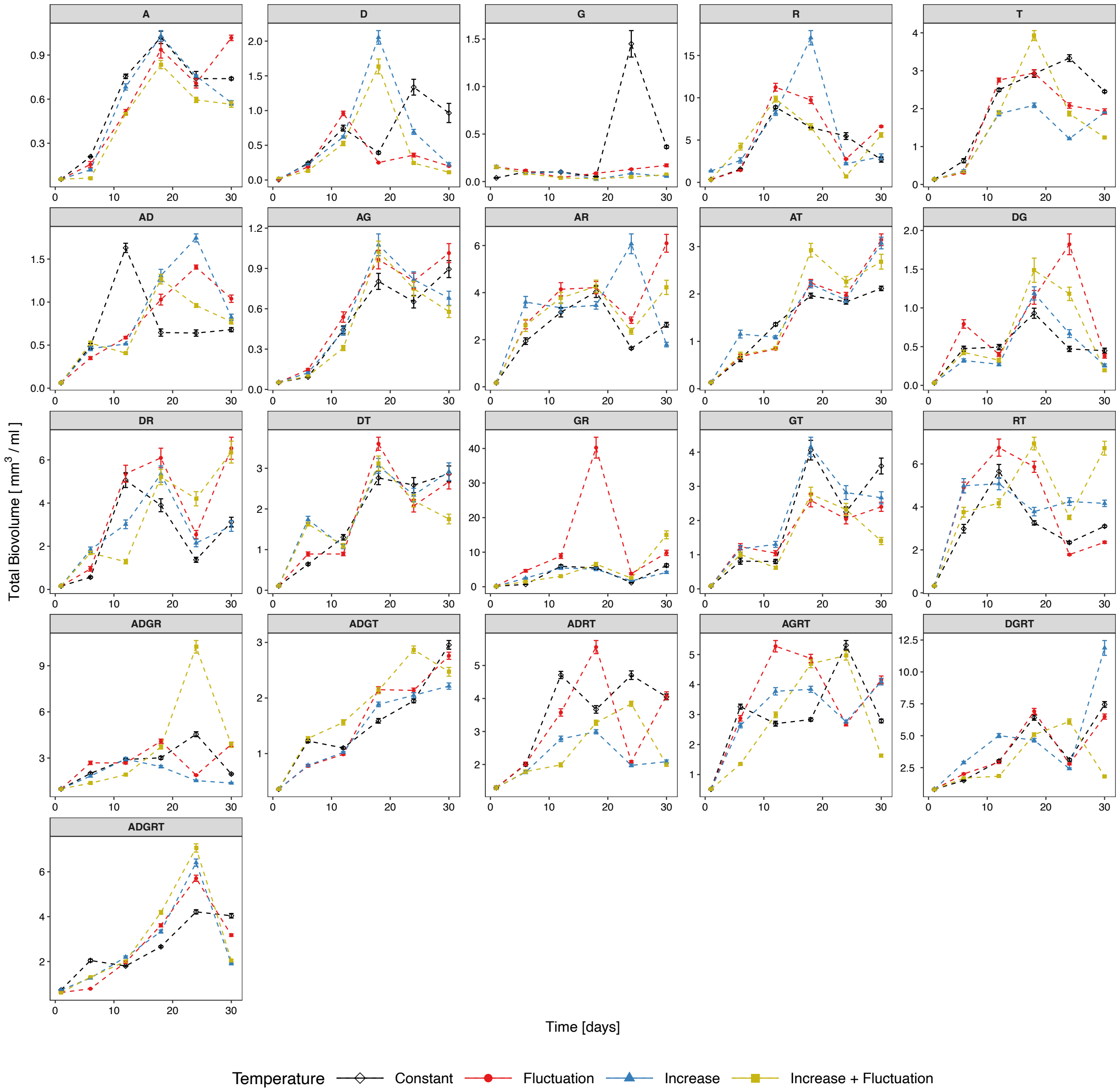


**Supporting Information Fig. S2.** Total biomass of species assemblages over time. Mean (± SE, *n* = 3) total biomass was measured from the total biovolume of the respective species compositions over time. Depending on species composition, temperature treatments frequently enhanced biomass production over time, except for most monocultures where temperature treatments decreased biomass compared to the constant temperature control. Species are abbreviated as A – *Asterionellopsis*, D – *Ditylum*, G – *Guinardia*, R – *Rhizosolenia*, T – *Thalassionema*, and species combinations are indicated by a combination of these abbreviations in each panel. Richness levels comprised species in monoculture (top row), two species (second and third row), four species (fourth row), and five species (bottom row).

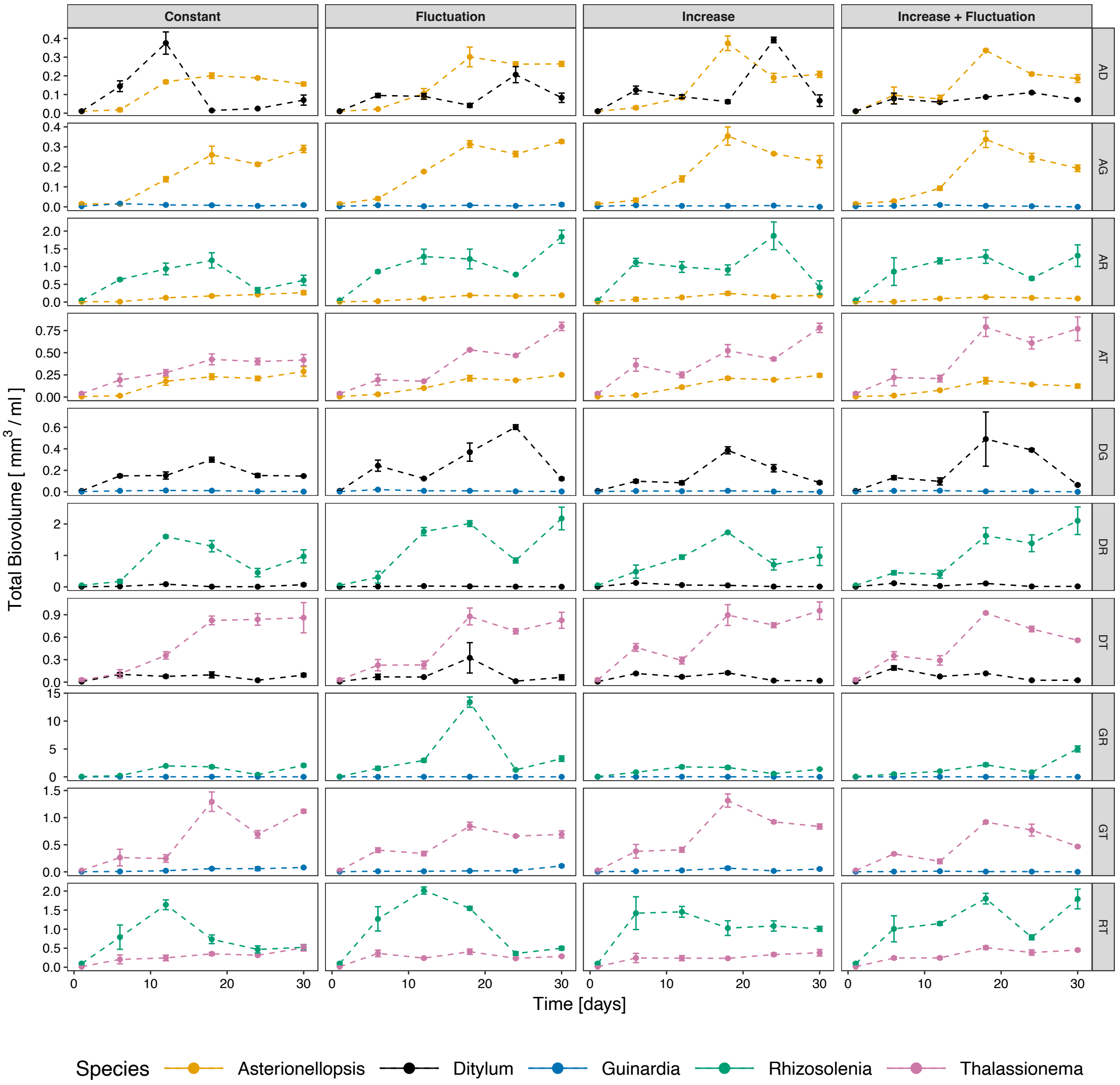


**Supporting Information Fig. S3.** Species-specific biomass of two species in mixtures over time. Mean (± SE, *n* = 3) total biovolume of each species in the two species combinations. Species in combinations are abbreviated as A – *Asterionellopsis*, D – *Ditylum*, G – *Guinardia*, R – *Rhizosolenia*, T – *Thalassionema*.

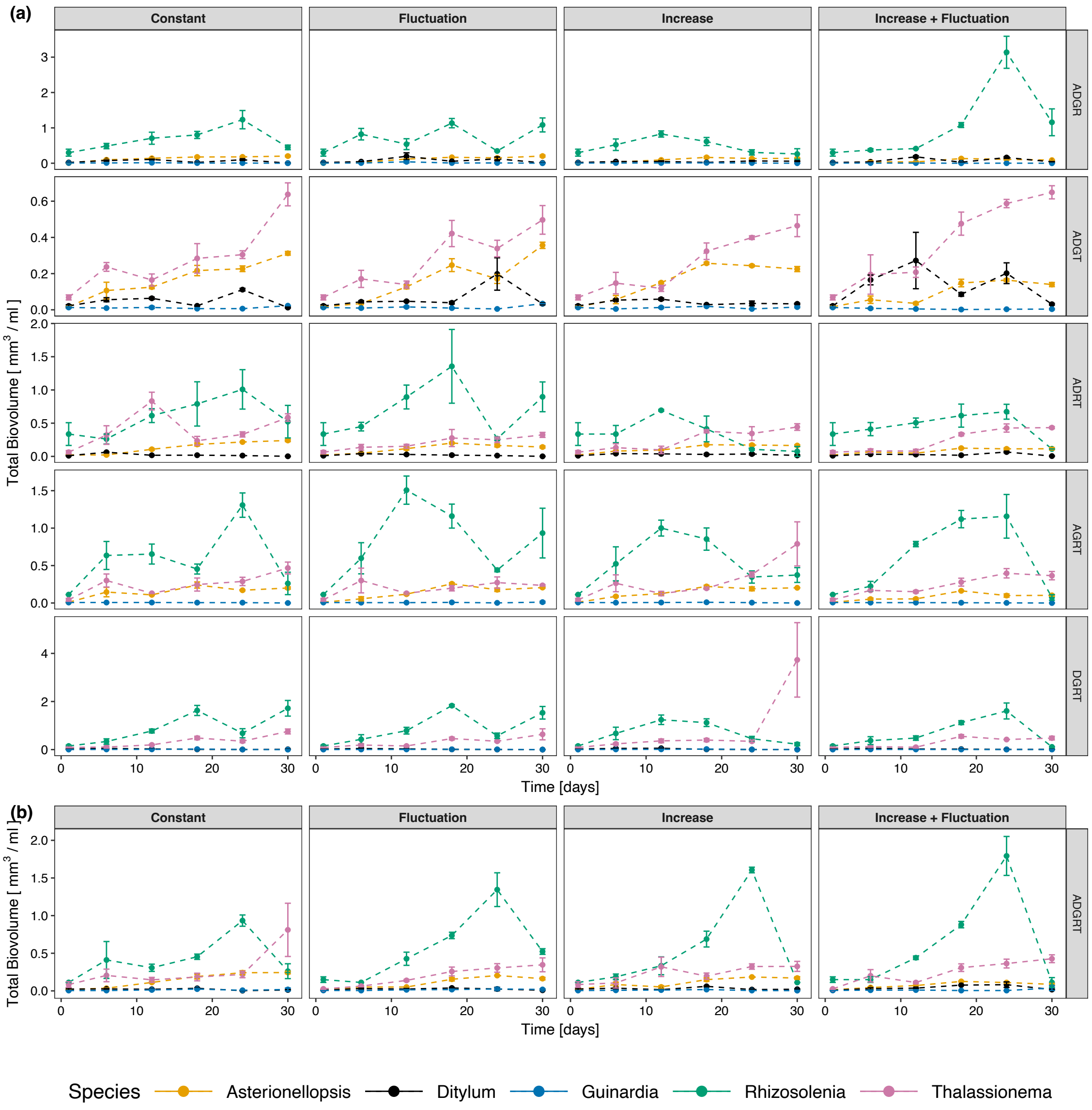


**Supporting Information Fig. S4.** Species-specific biomass in four-species and five-species assemblages over time. Mean (± SE, *n* = 3) total biovolume of each species in the four species (a) and five species assemblages (b). Species in combinations are abbreviated as A – *Asterionellopsis*, D – *Ditylum*, G – *Guinardia*, R – *Rhizosolenia*, T – *Thalassionema*.

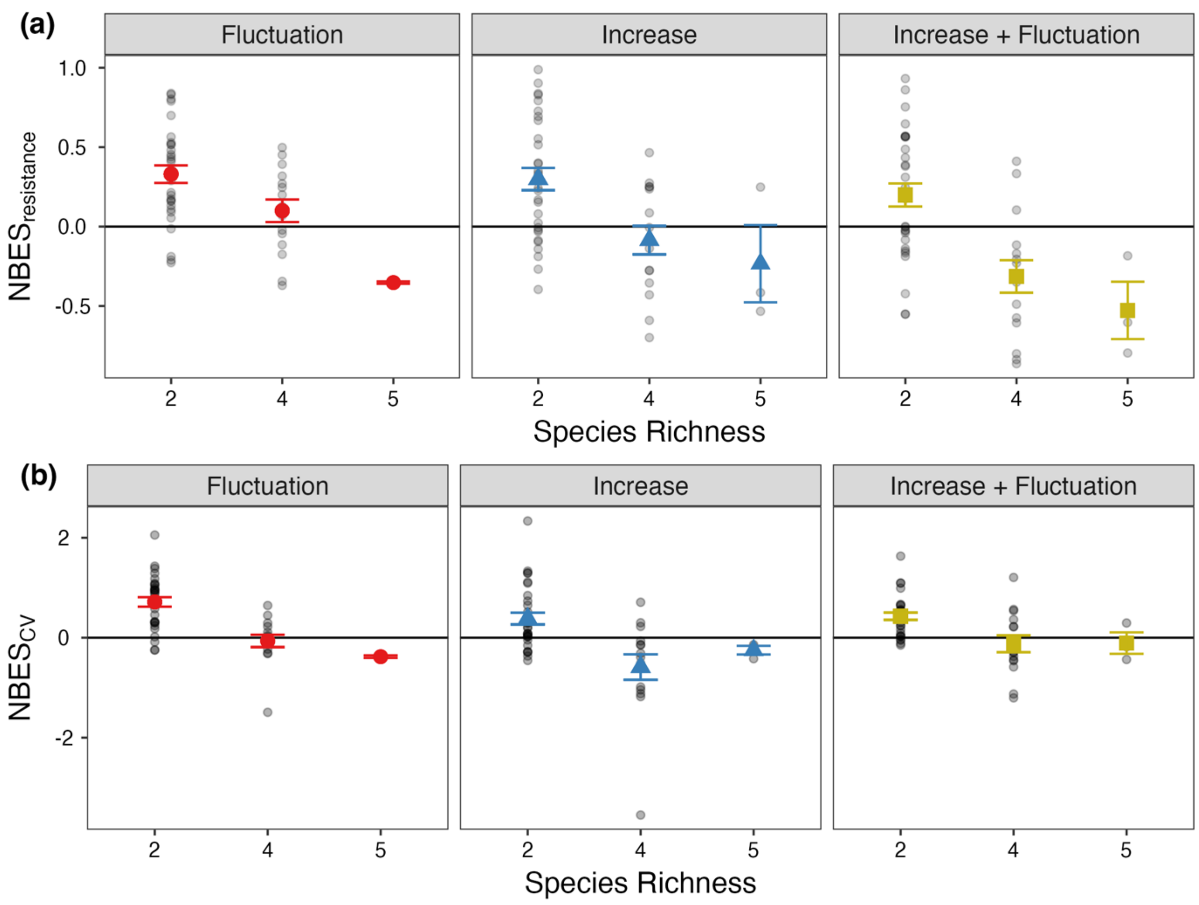


**Supporting Information Fig. S5.** The Net Biodiversity Effect on Stability for temporal variability and resistance. Mean ($n$ = 3 ± SE) net biodiversity effect on stability (NBES) based on resistance (a) and temporal variability (b) in our phytoplankton microcosms. Resistance was calculated as the difference in response ratios of multi-species assemblages compared to what is expected (a). Temporal variability was calculated as the difference in the coefficient of variation of response ratios of multi-species assemblages compared to what is expected from monoculture, over the species richness gradient (b).

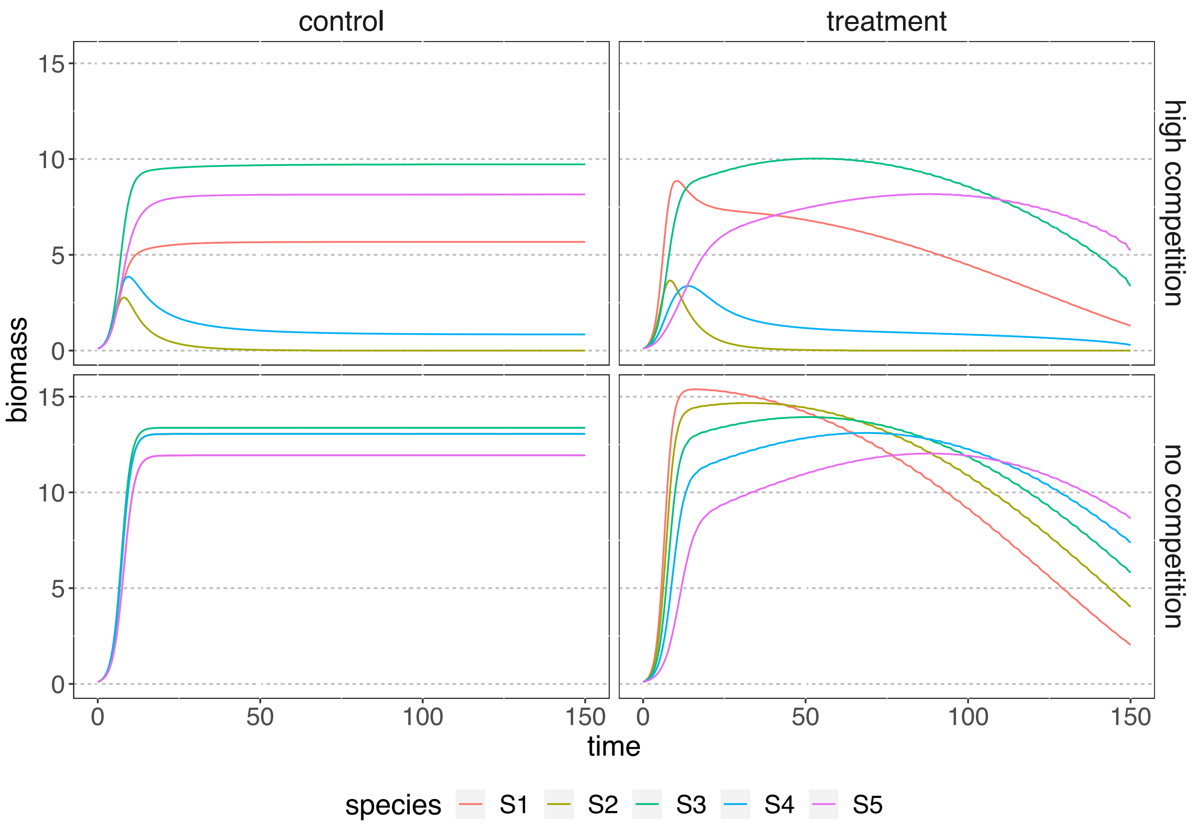


**Supporting Information Fig. S6**: Exemplary model runs of two communities where all species have different thermal optima and either strong competition or no competition under increasing temperatures and control conditions.
